# Total parenteral nutrition drives glucose metabolism disorders by modulating gut microbiota and its metabolites

**DOI:** 10.1101/2021.10.26.466009

**Authors:** Haifeng Sun, Peng Wang, Gulisudumu Maitiabula, Li Zhang, Jianbo Yang, Yupeng Zhang, Xuejin Gao, Jieshou Li, Bin Xue, Chao-Jun Li, Xinying Wang

## Abstract

The occurrence of glucose metabolism disorders is a potentially fatal complication of total parenteral nutrition (TPN). However, the mechanisms of TPN-associated glucose metabolism disorders remain unclarified. Given that the glucose metabolism was related to gut microbiome and TPN could induce the gut microbiota dysbiosis, we hypothesized that gut microbiota and its metabolites played the important roles in TPN-associated glucose metabolism disorders. By performing a cohort study of 256 type 2 IF patients given PN, we found that H-PN (PN>80%) patients exhibited insulin resistance and a higher risk of complications. Then, TPN and microbiome transfer mice model showed that TPN promoted glucose metabolism disorders by inducing gut microbiota dysbiosis; 16S rRNA sequencing demonstrated that the abundance of *Lactobacillacea*e was decreased in mice model and negatively correlated with HOMA-IR and lipopolysaccharide level in TPN patients. Untargeted metabolomics found that indole-3-acetic acid (IAA) was decreased in TPN mice, and the serum level was also decreased in H-PN patients. Furthermore, GLP-1 secretion regulated by IAA and aryl hydrocarbon receptor was reduced in TPN mice and patients; IAA or liraglutide completely prevented glucose metabolism disorders in TPN mice. In conclusion, TPN drives glucose metabolism disorders by inducing alteration of gut microbiota and its metabolites.

## Introduction

Intestinal failure (IF) is “the reduction of gut function below the minimum necessary for the absorption of macronutrients and/or water and electrolytes, such that intravenous supplementation is required to maintain health and/or growth”(1) and classified into acute (type 1), prolonged acute (type 2) or chronic (type 3).(1, 2) Type 2 IF persists for weeks/months and commonly arises after abdominal surgery for trauma, anastomotic leaks, vascular or viscous injury caused by another operation, enterocutaneous fistula, intestinal volvulus or mesenteric infarction.(3) Intermittent or continuous parenteral nutrition (PN) is a life-saving intervention for IF patients. (4) PN providing >80% of the patient’s energy requirement (H-PN) is defined as total parenteral nutrition (TPN) in IF patients.(5) TPN is associated with impairments in intestinal function,(6, 7) the gut barrier,(8) and mucosal immunity(9) which are thought to contribute to the development of adverse events such as infections and metabolic complications.(10) Hyperglycemia and hypoglycemia are the main metabolic complications of PN.(11) Hyperglycemia during PN can lead to immunosuppression and an increased susceptibility to enterogenic infection and is associated with various adverse outcomes including death.(12, 13) Hypoglycemia is also an acute and potentially fatal complication of TPN.(14, 15) Although the conventional theory suggests that PN-related glucose metabolism disorders is caused by excessive or insufficient infusion of dextrose and/or insulin, other mechanisms may also be involved.

There is evidence of a causal link between the gut microbiota and host metabolism,(16, 17) especially glucose handling.(18, 19) A recent study reported that TPN disrupted the gut microbiota and reduced microbiota-derived tryptophan metabolism.(20) The diverse repertoire of gut microbiota-derived metabolites plays a key role in the gut-multiorgan axis.(21) The gut microbiota produce numerous tryptophan metabolites, such as indole-3-acetic acid (IAA), tryptamine, indole-3-lactic acid and kynurenic acid. These metabolites bind to and activate the aryl hydrocarbon receptor (AhR).(22–24) Emerging evidence suggests that activation of AhR signaling improves insulin resistance and glucose handing.(25, 26) Interestingly, the gut microbiota and tryptophan metabolites also regulate the production of glucagon-like peptide-1 (GLP-1), which is an enteroendocrine hormone that regulates glucose homeostasis.(22, 27) However, the roles of microbiota-derived tryptophan metabolism and GLP-1 signaling in PN-related glucose metabolism disorders have not been elucidated.

The aim of present study was to determine the prevalence of glucose metabolism disorders in type 2 IF patients given PN, investigate the role of the gut microbiota and its metabolites in the development of TPN-associated glucose metabolism disorders and elucidate the underlying mechanisms.

## Results

### A higher level of energy provision by PN is associated with glucose metabolism disorders and clinical complications in patients with type 2 IF

Metabolic status and clinical complications were evaluated in 256 type 2 IF patients who required continuous or intermittent PN for several weeks or months. The PN regimens and clinical characteristics of the patients are presented in Table S2 and Table S3, respectively. Short bowel syndrome and ileus were more common in patients given H-PN (PN providing >80% of total energy) than in those given L-PN (PN providing ≤80% of total energy), whereas extensive parenchymal disease was a less common underlying disease in the H-PN group (P<0.05,Table S2). Additionally, the H-PN group had a higher body mass index (BMI), longer hospital stay, higher hospitalization cost, and poorer nutritional status according to the patient-generated subjective global assessment (PG-SGA) scale than patients in the L-PN group (P<0.05; Table S3). However, the residual small intestinal length in 75 patients with SBS has no statistical difference between the two groups (71.6±48.1cm VS 69.9±42.5 cm P=0.06, Table S2). Notably, the H-PN group had higher levels of fasting blood glucose (6.27±0.16 vs. 5.14±0.09 mmol/L, P<0.0001; Fig. 1A), fasting blood insulin (10.57±0.58 vs. 6.33±0.36 mIU/L, P<0.0001; Fig. 1B) and homeostasis model assessment of insulin resistance (HOMA-IR; 3.23±0.19 vs. 1.37±0.09, P<0.0001; Fig. 1C) than the L-PN group, whereas there was no significant difference between the two groups in total energy intake (27.98±0.88 vs. 29.75±0.56 kcal/kg/d, P=0.0774; Fig. 1D). Furthermore, the degree of insulin resistance (HOMA-IR) was strongly positively correlated with the proportion of energy provided by PN (r=0.6019, P<0.0001; Fig. 1E) and negatively correlated with skeletal muscle mass (r=-0.3899, P<0.001; Fig. 1F), fat-free mass (r=−0.4104, P<0.001; Fig. 1G) and body protein mass (r=-0.4146, P<0.001; Fig. 1H) expressed as a proportion of total body weight. These data suggest that insulin resistance in IF patients is associated not only with body composition but also with the proportion of energy provided by PN.

**Figure1.**
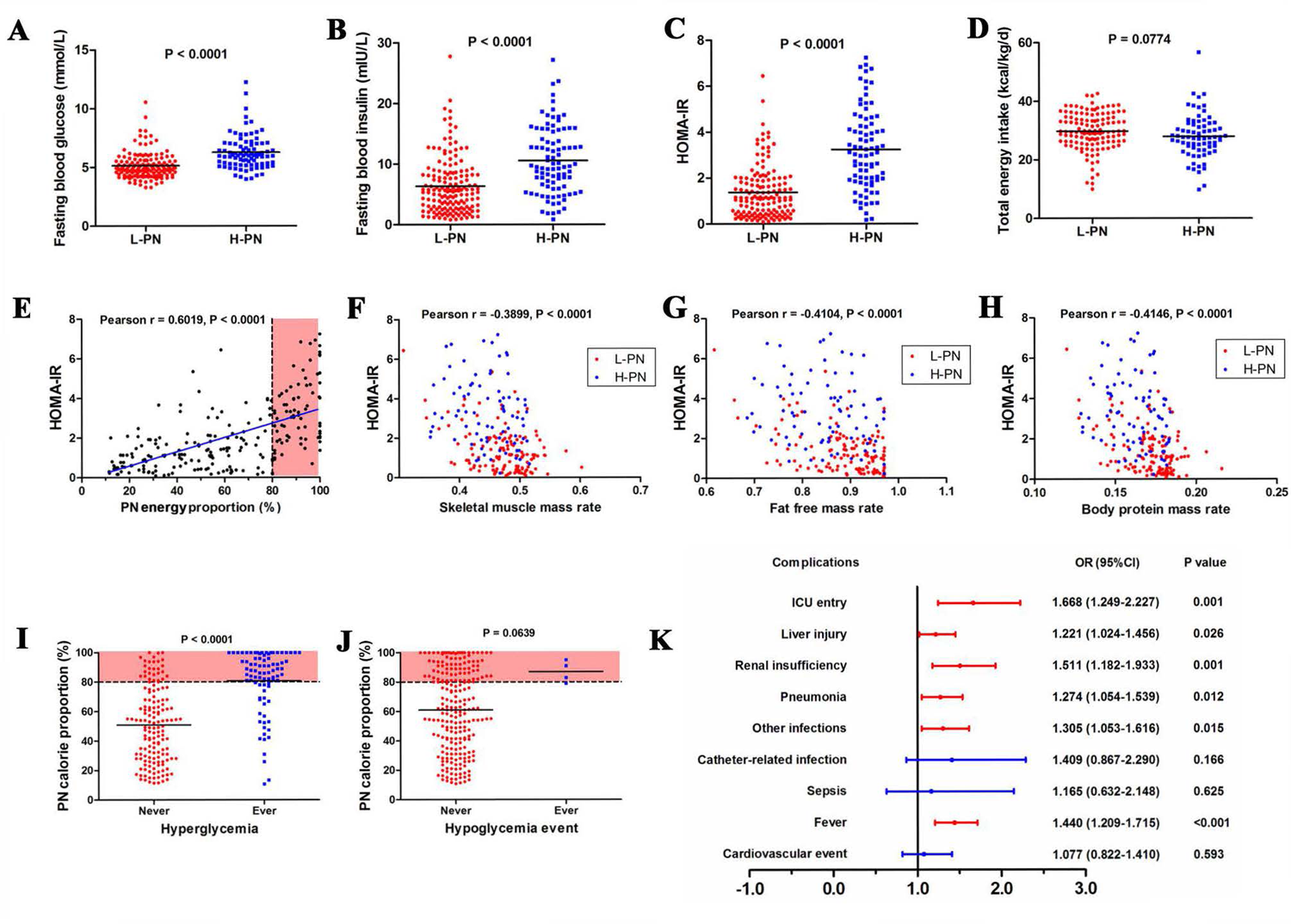
A higher level of energy provision by parenteral nutrition (PN) is associated with glucose metabolism disorders and clinical complications in patients with type 2 intestinal failure (IF). A-D. The average levels of fasting blood glucose (A), fasting blood insulin (B), homeostasis model assessment of insulin resistance (HOMA-IR) (C) and daily energy intake (D) in hospitalized patients with IF given PN. L-PN: ≤80% of energy provided by PN; H-PN: >80% of energy provided by PN. E-H. Correlation between HOMA-IR value and the proportion of energy provided by PN (E), skeletal muscle mass relative to total body mass (F), fat-free mass relative to total body mass (G) and body protein mass relative to total body mass (H) in patients with IF. I-J. The relationship between the proportion of energy provided by PN and the occurrence of hyperglycemic events (I) and hypoglycemic events (J) during hospitalization. K. Forest plot showing the impact of HOMA-IR value on clinical complications in patients with IF.

Analysis of outcomes during hospitalization revealed that hyperglycemia (Fig. 1I) and hypoglycemic events (Fig. 1J) occurred predominantly in the H-PN group. Moreover, HOMA-IR was associated with increased risks of intensive care unit (ICU) transfer (OR [odds ratio]=1.668, P=0.001), liver injury (OR=1.221, P=0.026), renal insufficiency (OR=1.511, P=0.001), pneumonia (OR=1.274, P=0.012), other infections (OR=1.305, P=0.015) and fever (OR=1.440, P<0.001) during hospitalization but not catheter-related infection, sepsis or cardiovascular events (Fig. 1K). The above observations indicate that the development of insulin resistance in type 2 IF patients receiving PN increases the risks of unfavorable outcomes.

### TPN impairs insulin sensitivity and liver glycogen deposition in mice

The possible association between PN and insulin sensitivity was further explored in a mouse model. IPGTTs (Figs. 2A and S2A) and IPITTs (Figs. 2B and S2B) showed that insulin sensitivity was impaired in mice given TPN. Although there was no significant difference in fasting glucose level between groups (Fig. 2C), fasting insulin level (Fig. 2D) and HOMA-IR (Fig. 2E) were significantly higher in mice given TPN than in control mice fed a standard chow diet, which suggests that TPN might result in insulin resistance. Glycogen deposition in hepatocytes was lower in the TPN group than in the control group (Fig. 2F and 2G). Furthermore, insulin-induced phosphorylation of insulin receptor substrate-1 (IRS1), Akt and glycogen synthase kinase-3β (GSK3β) in hepatocytes was lower in the TPN group than in the control group, indicating that TPN impaired the liver’s response to insulin (Figs. 2H). The above observations raise the possibility that reduced insulin sensitivity may contribute to TPN-associated glucose metabolism disorders (Fig. S2C). Additionally, alanine transaminase (ALT) and aspartate transaminase (AST) were significantly higher in mice given TPN than in controls (Fig. 2I and 2J), suggesting hepatic dysfunction. Taken together with the clinical data, these results suggest that TPN induces insulin resistance and impaired hepatic function that aggravates glucose metabolism disorders.

**Figure 2.**
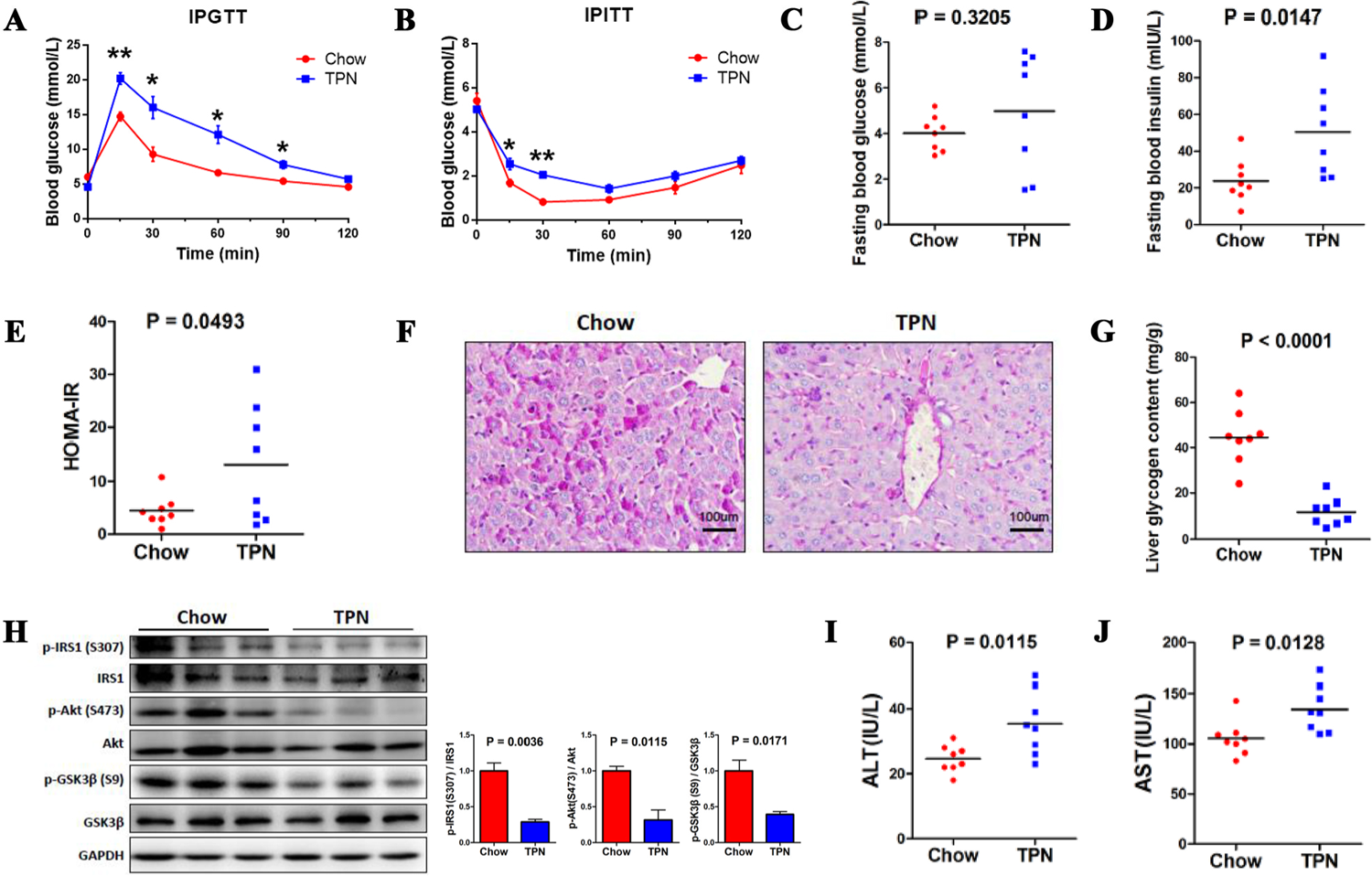
Total parenteral nutrition impairs insulin sensitivity, liver glycogen deposition, insulin-dependent signaling and hepatic function in mice (n=8 each group, 3 times repeat). A-E. Intraperitoneal glucose tolerance test (A), intraperitoneal insulin tolerance test (B), fasting blood glucose level (C), fasting blood insulin level (D) and homeostasis model assessment of insulin resistance (HOMA-IR) value (E). * P<0.05, ** P<0.01, TPN group vs. Chow group. F-G. Representative histologic images showing deposition of glycogen in mouse hepatocytes (F) and quantification of the average levels of liver glycogen (G). H. Western blot (Left) and semiquantitative analyses (Right) of insulin-driven glycogen synthesis signaling (IRS1-Akt-GSK3) in the liver. I-J. Serum alanine transaminase (ALT) and aspartate transaminase (AST) concentrations.

### Alterations in the composition of the gut microbiota contribute to TPN-induced insulin resistance and hepatic glycogen deficit

16S sequencing revealed that TPN altered the gut microbiome profiles (Fig. 3A). Specifically, TPN did not affect bacterial diversity (Fig. 3B) but significantly altered the microbiota composition (Fig. 3C and 3D). Heatmaps showing microbial composition at the phylum and genus levels are shown in Fig. S3A,B. TPN significantly decreased the relative abundances of *Lactobacillaceae* (P=0.0299) and *Rhodocyclaceae* (P=0.0336) and increased the relative abundances of *Sphingomonadaceae* (P=0.0074) and *Ruminococcaceae* (P=0.0115; Fig. 3E).

**Figure 3.**
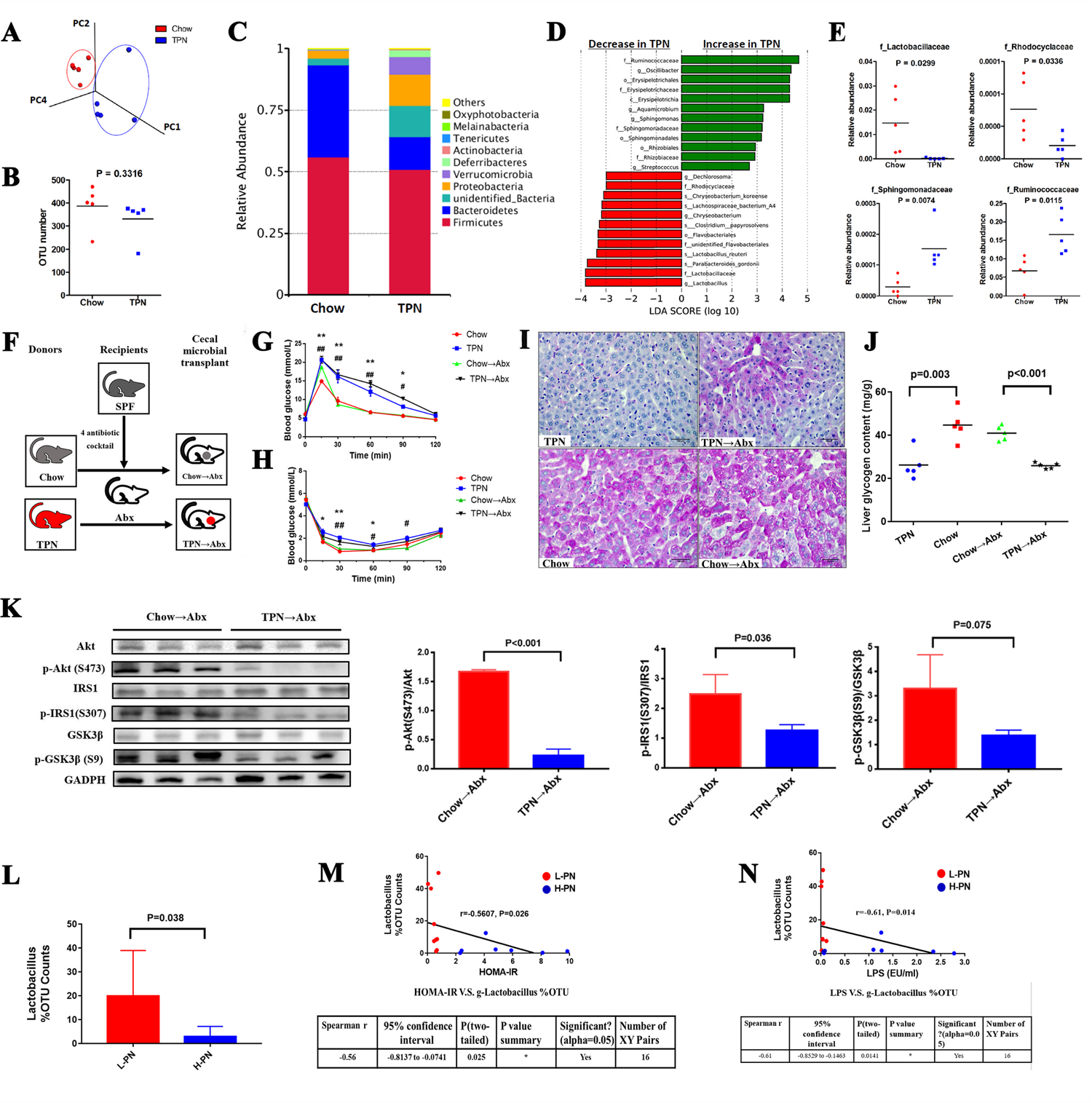
Total parenteral nutrition (TPN) alters the composition of the gut microbiota (n=5 each group, 3 times repeat). A. Principal coordinate analysis plot of Bray-Curtis distances. B. Bacterial diversity analysis based on operational taxonomic units (OTUs). C. Bacterial proportions at the phylum level. D. Bacterial discriminant analysis based on the linear discriminant analysis (LDA) score. E. Relative abundance of significantly different bacteria between the Chow and TPN groups. F. Schematic of the fecal microbiota transplantation experiments. G-H. Intraperitoneal glucose tolerance test (G) and intraperitoneal insulin tolerance test (H) in the Chow---+Abx and TPN---+Abx groups of mice.* P<0.05, ** P<0.01, TPN group vs. Chow group; # P<0.05, ## P<0.01, Chow---+Abx group vs TPN---+Abx group. I-J. Representative histologic images showing deposition of glycogen in mouse hepatocytes (I) and quantification of the average levels of liver glycogen (J). K. Western blot (Left) and semiquantitative analyses (Right) of insulin-driven glycogen synthesis signaling (IRS1-Akt-GSK3) in the liver. L. Lactobacillus abundance in patients with intestinal failure. L-PN: ≤80% of energy

Next, we performed cecal microbiota transplantation to explore whether alterations in the microbiota contributed to TPN-associated insulin resistance and liver glycogen deficit. The recipient (Abx) mice were pretreated for 2 weeks with antibiotics (ampicillin, neomycin, metronidazole and vancomycin), which depleted >95% of the intestinal microbiota according to quantitative polymerase chain reaction (qPCR) analysis of the fold-change in 16S rDNA (Fig. S3C). Cecal microbiota collected from mice in the Chow (control) and TPN groups (at 7 days) were given by gavage to Abx mice (Chow→Abx and TPN→Abx groups; Fig. 3F). When compared with the Chow and Chow→Abx groups, the TPN and TPN→Abx groups exhibited decreased glucose tolerance (Figs. 3G and S3D), insulin resistance (Figs. 3H and S3E) and hepatic glycogen deficit (Fig. 3I and 3j). Additionally, IRS1, Akt and GSK3β phosphorylation was lower in TPN→Abx mice than in Chow→Abx mice (Figs. 3K), suggesting a reduction in insulin signaling.

The effects of PN on the intestinal flora were further examined in 16 patients with IF (recruited separately from the main cohort of 256 patients). The clinical characteristics of these 16 patients are shown in Table S4. 16S RNA sequencing of fecal or jejunostomy liquid samples revealed that *Lactobacillus* abundance was lower in patients given H-PN than in patients given L-PN (P=0.038; Fig. 3L). Furthermore, *Lactobacillus* abundance was negatively correlated with HOMA-IR (Fig. 3M) and serum lipopolysaccharide (LPS) concentration (Fig. 3N).

### TPN induces insulin resistance by reducing the levels of tryptophan metabolites

Examination of the cecal contents of mice demonstrated differences in metabolite profiles between the TPN and control groups (Fig. 4A). IAA, a product of microbial tryptophan metabolism, was an outlying downregulated metabolite in the TPN group, while docosahexaenoic acid (DHA) was an outlying upregulated metabolite in the TPN group (Fig. 4B). Correlation analysis was performed to determine the potential associations of the TPN-altered microbes and metabolites. The correlation heatmap revealed that TPN-induced depletion of *Lactobacillaceae* and *Muribaculum* was positively correlated with IAA, indole-3-lactic acid (ILA), indole and kynurenic acid, while TPN-induced enrichment of *Akkermansia, Helicobacter* and *Oscilibacter* was negatively associated with indole metabolites (Fig. 4C). The metabolite heatmap is shown in Fig. S4A. Additionally, Kyoto Encyclopedia of Genes and Genomes (KEGG) analysis indicated that tryptophan metabolism was the second most concentrated differential metabolic pathway between the two groups (Fig. S4B).

**Figure 4.**
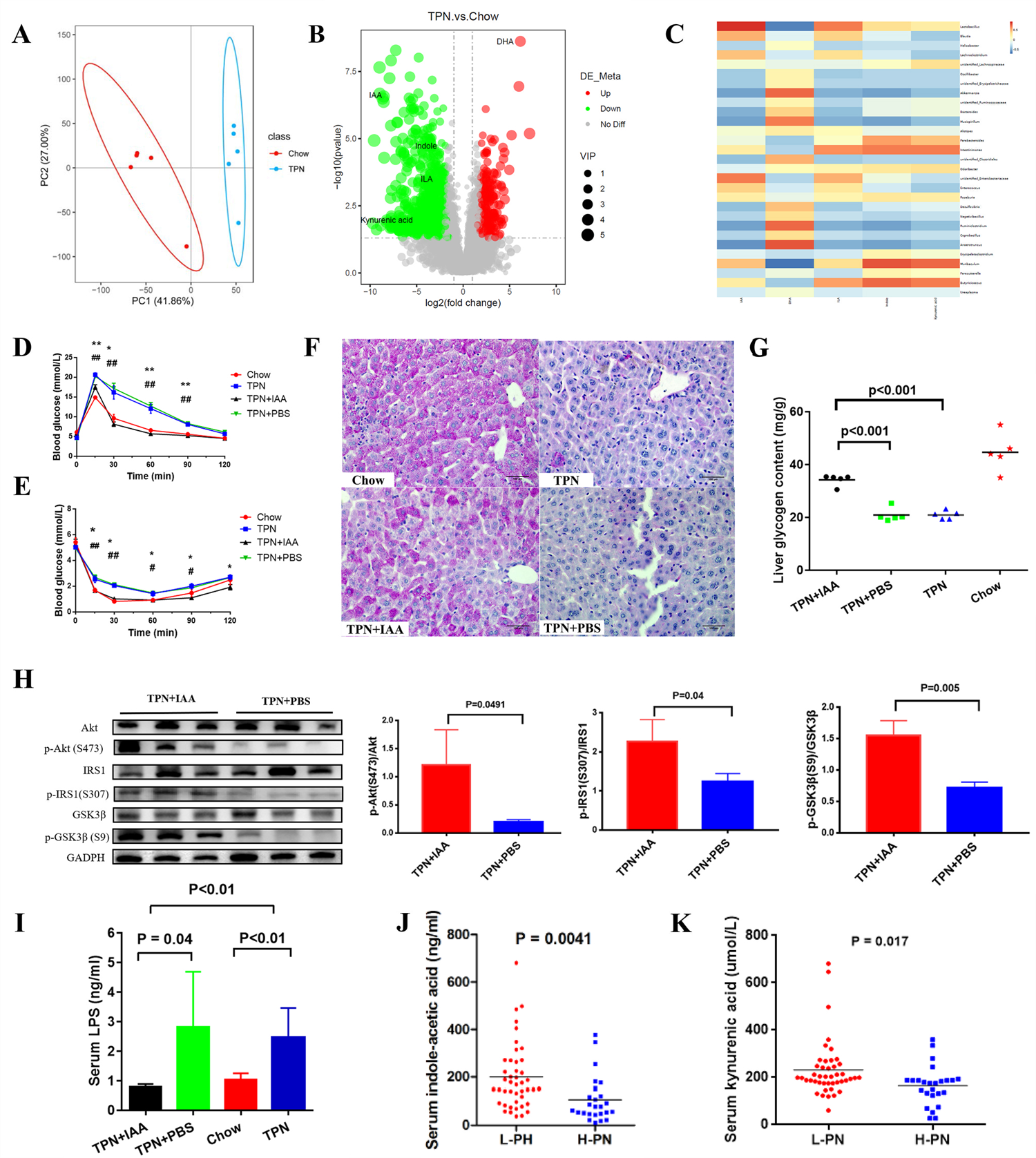
Total parenteral nutrition (TPN) promotes poor insulin sensitivity by reducing the levels of tryptophan metabolites including indole-3-acetic acid (IAA) (n=5 each group, 3 times repeat). A. Principal component analysis based on metabolite composition. B. Volcano plot identifying differential serum metabolites between mice in the TPN and Chow groups. The outlying metabolites are indicated. C. Correlation analysis showing the associations between TPN-altered microbes and metabolites. D-E. Intraperitoneal glucose tolerance tests (D) and intraperitoneal insulin tolerance tests (E) for the four groups of mice.* P<0.05, ** P<0.01, TPN group vs. Chow group;# P<0.05, ## P<0.01, TPN+IAA group vs TPN+PBS group. F-G. Representative histologic images showing deposition of glycogen in mouse hepatocytes (F) and quantification of the average levels of liver glycogen (G). H. Western blot (Left) and semiquantitative analyses (Right) of insulin-driven glycogen synthesis signaling (IRSl-Akt-GSK3) in the liver. I. Serum concentration of lipopolysaccharide (LPS). J-K. Serum concentrations of IAA (J) and kynurenic acid (K) in patients with intestinal failure. L-PN: ≤80% of energy provided by PN; H-PN: >80% of energy provided by PN.

Therefore, we explored whether supplementation with IAA would improve glucose tolerance and insulin sensitivity in TPN mice. IPGTTs (Fig. 4D and S4C) and IPITTs (Figs. 4E and S4D) showed that IAA administration enhanced glucose tolerance and improved insulin sensitivity in TPN mice. Periodic acid-Schiff (PAS) staining showed that intraperitoneal injection of IAA significantly increased liver glycogen content in the TPN group to levels comparable to those of chow-fed mice (Fig. 4F,G). Moreover, IAA administration rescued the phosphorylation of IRS1, Akt and GSK3β (Fig. 4H) and reduced the circulating level of LPS in TPN mice (Fig. 4I).

Plasma concentrations of IAA and kyunrenic acid were measured in 69 of the 256 IF patients given PN to assess whether tryptophan metabolite levels were influenced by the energy provision level. The plasma levels of IAA and kynurenic acid were significantly lower in the H-PN group than in the L-PN group (P=0.004 and P=0.017, respectively; Fig. 4J,K). Taken together, these data suggest that TPN reduces the production of IAA and other tryptophan metabolites (such as kynurenic acid) by the intestinal flora and that this may play an important role in TPN-associated glucose metabolism disorders.

### TPN-related insulin resistance is associated with inactivation of indole/AhR signaling

Next, we performed experiments in mice using an AhR inhibitor (CH223191) to examine whether indole/AhR signaling is involved in TPN-induced insulin resistance. Chow-fed mice administered CH223191 exhibited insulin resistance and reduced liver glycogen deposition (similar to TPN mice) that were not improved by IAA injection (Fig. 5A–D and Fig. S5A,B). CH223191 also suppressed GSK3β and Akt phosphorylation in chow-fed mice, and these effects were not reversed by IAA (Fig. 5E).

**Figure 5.**
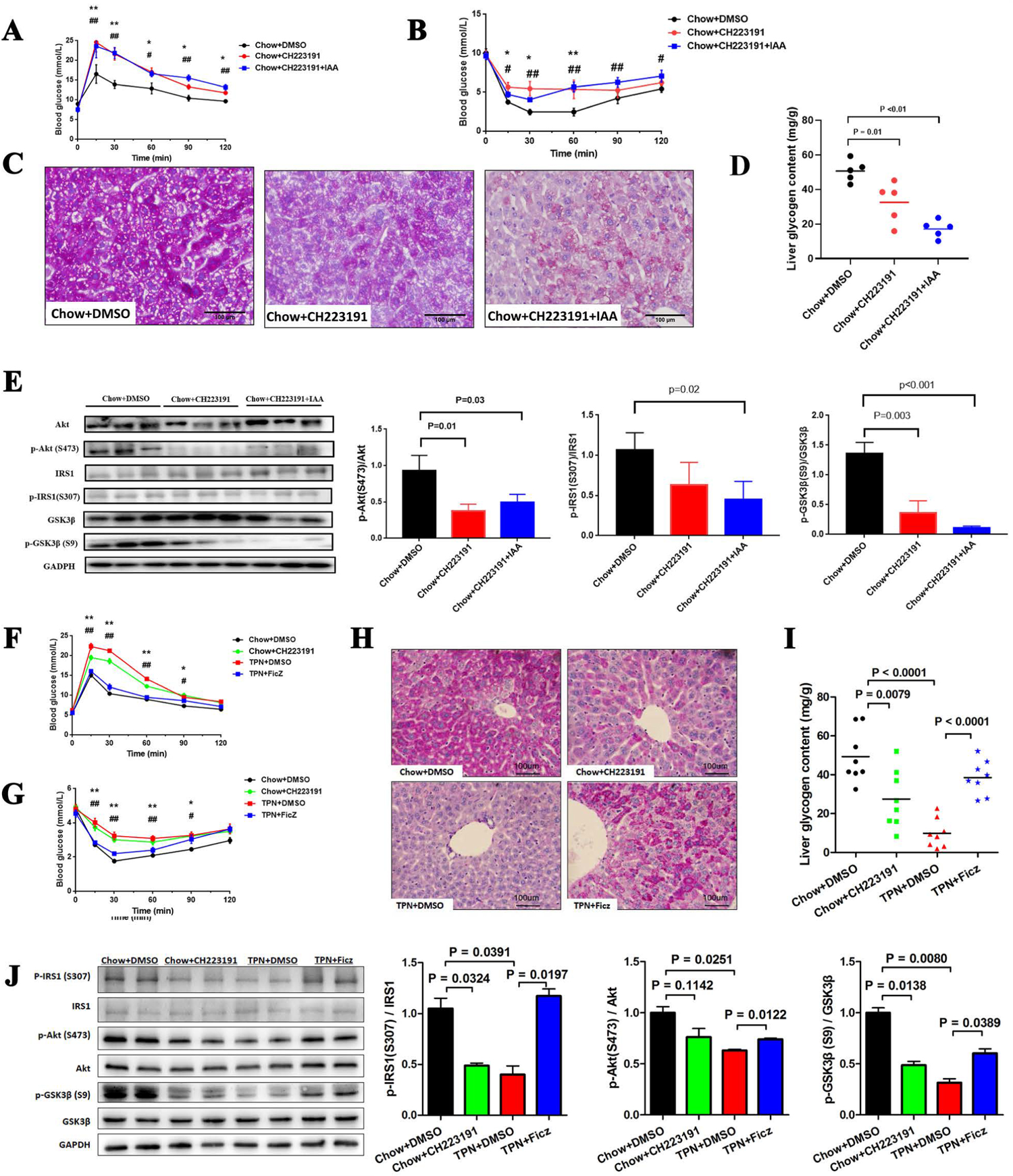
Inactivation of indole/aryl hydrocarbon receptor signaling contributes to parenteral nutrition-related impairment of insulin sensitivity. A-B. Intraperitoneal glucose tolerance tests (A) and intraperitoneal insulin tolerance tests (B).(* P<0.05, ** P<0.01, Chow+DMSO group vs. Chow+CH223191 group;# P<0.05, ## P<0.01, Chow+CH223191 group vs Chow+CH22319+IAA group.) C-D. Representative histologic images showing deposition of glycogen in mouse hepatocytes (C) and quantification of the average levels of liver glycogen (D) E. Western blot (Left) and semiquantitative analyses (Right) of insulin-driven glycogen synthesis signaling (IRS1-Akt-GSK3) in mouse liver. (n=5 each group, 3 times repeat). F-G. Intraperitoneal glucose tolerance tests (F) and intraperitoneal insulin tolerance tests (G). *P<0.05, ** P<0.01, Chow+DMSO group vs. Chow+CH223191 group;# P<0.05, ## P<0.01, TPN+DMSO group vs TPN+Ficz group. H-I. Representative histologic images showing deposition of glycogen in mouse hepatocytes (H) and quantification of the average levels of liver glycogen (I). J. Western blot (Left) and semiquantitative ana1 ses (Right) of insulin-driven glycogen synthesis signaling (IRS1-Akt-GSK3) in the liver (n=8 each group, 3 times repeat).

Additional experiments were performed using an AhR agonist (Ficz). IPGTTs (Figs. 5F and S5C) and IPITTs (Figs. 5G and S5D) showed that activation of AhR signaling by Ficz improved glucose tolerance and insulin sensitivity in TPN mice. Ficz treatment also enhanced hepatic glycogen deposition (Fig. 5H and 5I) and recovered the phosphorylation of IRS1, Akt and GSK3β (Fig. 5J) in TPN mice. These data suggest that TPN-induced insulin resistance and liver dysfunction are associated with inactivation of AhR signaling due to reductions in the levels of indole derivatives.

### Inactivation of indole/AhR signaling inhibits GLP-1 production to induce metabolic complications in TPN mice

Glucose metabolism disorders and liver steatosis due to dietary or genetic causes are reversed by indole production from tryptophan, AhR activation, secretion of GLP-1 and improved intestinal barrier function.[25] Measurements made in 69 of the 256 patients revealed that the plasma GLP-1 level was significantly lower in H-PN patients than in L-PN patients (Fig. 6A) despite no difference between groups in the length of the residual small bowel. Experiments using chow-fed mice demonstrated that inhibition of indole/AhR signaling with CH223191 induced a significant decrease in the plasma GLP-1 level, while activation of indole/AhR signaling with Ficz increased GLP-l levels in TPN mice (Fig. 6B). Since GLP-1 is synthesized and released from L cells predominantly in the terminal ileum and colon, we evaluated GLP-1 expression in different segments of the gut from chow- and TPN-fed mice. Immunostaining experiments revealed that the number of GLP-1-positive cells in both the terminal ileum and colon was decreased by CH223191 in chow-fed mice and increased by Ficz in TPN-fed mice (Fig. 6C). The mRNA level of proglucagon (the precursor of GLP-1; Fig. S6A–B) and the protein level of GLP-1 (Fig. S6C) in the terminal ileum and colon were also reduced by CH223191 in chow-fed mice and enhanced by Ficz in TPN-fed mice. These data suggest that indole/AhR signaling plays a key role in the regulation of GLP-1 production. Hence, this pathway might be a suitable target for the treatment of TPN-related metabolic disturbances such as insulin resistance and liver dysfunction.

**Figure 6.**
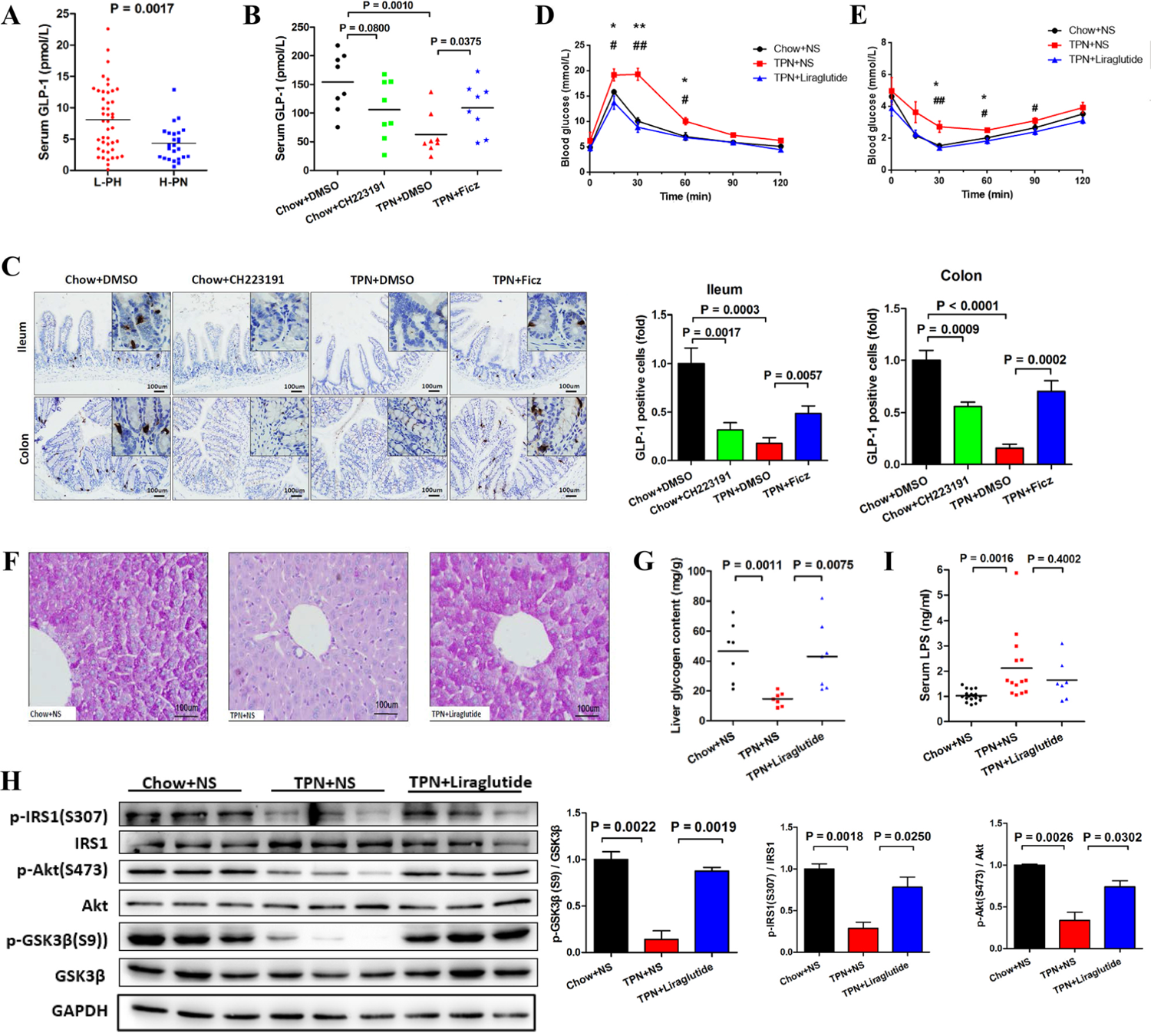
Inactivation of indole/aryl hydrocarbon receptor signaling reduces glucagon-like peptide-I production to cause parenteral nutrition (PN)-related impairment of insulin sensitivity. (n=8 each group, 3 times repeat) A. Serum glucagon-like peptide-I (GLP-I) concentration in patients with intestinal failure. L-PN: ≤80% of energy provided by PN; H-PN: >80% of energy provided by PN. B. Serum GLP-I concentration in the four groups of mice. C. Immunostaining of GLP-I in mouse terminal ileum and colon. D-E. Intraperitoneal glucose tolerance tests (D) and intraperitoneal insulin tolerance tests (E). * P<0.05, ** P<0.01, Chow+NS group vs. TPN+NS group;# P<0.05, ## P<0.01, TPN+NS group vs TPN+Liraglutide group. F-G. Representative histologic images showing deposition of glycogen in mouse hepatocytes (F) and quantification of the average levels of liver glycogen (G). H. Western blot (Left) and semiquantitative analyses (Right) of insulin-driven glycogen synthesis signaling (IRSI-Akt-GSK3) in the liver. I. Serum lipopolysaccharide (LPS) concentration.

The relationship between decreased GLP-1 production and the metabolic complications of TPN were further investigated in mice administered liraglutide, a GLP-1 analog used clinically to treat diabetes mellitus. Liraglutide significantly improved glucose clearance (Figs. 6D and S6D), insulin sensitivity (Figs. 6E and S6E) and hepatic glycogen deposition (Fig. 6F and 6G) in the TPN group. Additionally, liraglutide rescued the phosphorylation of IRS1, Akt and GSK3β in TPN mice (Figs. 6H). Liraglutide also augmented the body composition of TPN mice as evidenced by a trend toward an increase in the gastrocnemius/body weight ratio (P=0.0998; Fig. S6F) and a significant decrease in the epididymal fat/body weight ratio (P=0.0121; Fig. S6G). Although liraglutide treatment was associated with numerical decreases in ALT (79.80±11.67 vs. 56.63±7.35 IU/L) and AST (150.6±12.50 vs. 122.3±10.88 IU/L), the apparent reductions were not statistically significant (Fig. S6H). Liraglutide administration also had no significant effect on liver edema (Fig. S6I) or plasma LPS level (Fig. 6I). Finally, despite an apparent trend toward a reduction in 7-day mortality in TPN-fed mice after liraglutide treatment, there was no significant difference between the liraglutide-treated and control groups (12.5% vs. 41.7%; Fig. S6J).

## Discussion

PN is an essential and life-saving treatment for IF,(28) but complications such as metabolic disorders and infection can seriously affect patients’ quality of life.(10) The present study provides new insights into the mechanism of TPN-induced metabolic complications and identifies potential targets for therapeutic interventions to prevent these complications. We have shown that type 2 IF patients given TPN exhibited severe insulin resistance that was associated with adverse complications. Furthermore, we have demonstrated for the first time that TPN-induced glucose metabolism disorders were associated with gut microbiota dysbiosis. Specifically, our findings indicate that TPN can induce insulin resistance by altering the composition of the gut microbiota and reducing the levels of metabolites such as IAA, which in turn decreases AhR-mediated signaling and GLP-1 production.

Conventional theory suggests that improper infusion of dextrose contributes to TPN-related glucose metabolism disorders, i.e., excessive dextrose infusion during TPN leads to hyperglycemia, whereas suddenly weaning off PN and compensatory hyperinsulinemia result in hypoglycemia.(29, 30) However, we found that TPN was associated with insulin resistance in both patients and mice. Moreover, TPN suppressed glycogen deposition in mice, in agreement with previous reports indicating that TPN reduced liver glycogen composition after 4 days.(31, 32) We propose that TPN-related glucose metabolism disorders results not only from improper intravenous infusion of dextrose but also from the development of insulin resistance that impairs glucose utilization and leads to hyperglycemia. Inadequate glycogen storage compromises the body’s ability to maintain blood glucose stability when PN is suddenly removed, which predisposes to hypoglycemia. Thus, we propose that TPN can impair insulin sensitivity and induce glucose metabolism disorders.

Accumulating evidence suggests that the human gut microbiota can influence various pathologic conditions including metabolic disorders.(16–19, 33) We therefore explored the potential involvement of the gut microbiota in TPN-induced glucose metabolism disorders in mice. We found that TPN was associated with decreased abundance of *Lactobacillaceae* and *Rhodocyclaceae* and increased abundance of *Sphingomonadaceae* and *Ruminococcaceae.* The findings are consistent with published studies describing TPN-associated changes in the gut microbiota. For example, Heneghan et al. reported that TPN led to a reduction in Firmicutes (which includes *Lactobacillaceae*) and an increase in Bacteroidetes and Proteobacteria in mice.(34) Interestingly, our study showed that transplantation of cecal contents induced glucose metabolism disorders in recipient mice and that the reduction in *Lactobacillaceae* was associated with a decrease in microbiota-derived tryptophan metabolism. Wang et al. also detected a decrease in the abundance of tryptophan metabolites.(20) These findings agree well with the present study and raise the possibility that alterations in tryptophan metabolite levels due to disruption of the gut flora might play a key role in PN-related complications such as insulin resistance.

Bacteria-derived metabolites such as short chain fatty acids,(35) serotonin(36) and indole(25) are believed to effect host-microbial cross-talk and the development of metabolic diseases. Among the metabolites generated by the gut microbiota, indole and its derivatives have been reported to have various metabolic benefits which they bring about by binding to and activating the AhR.(22) Natividad et al. concluded that metabolic syndrome in mice and humans is associated with a reduced ability of the gut microbiota to produce tryptophan metabolites that activate the AhR, and the authors showed that supplementation with a *Lactobacillus* strain or an AhR agonist ameliorated the metabolic impairments.(25) We found that the levels of several tryptophan metabolites, including IAA, indole-3-lactic acid, indole acetate, 5-hydroxyindole, 5-hydroxyindole-3-acetic acid and indole carbaldehyde, were significantly lower in TPN mice than in chow-fed mice. Furthermore, the plasma IAA concentration was significantly lower in H-PN patients than in those given L-PN. Notably, supplementation with IAA or administration of an AhR agonist (Ficz) improved glucose tolerance, insulin sensitivity, liver glycogen content and the phosphorylation of IRS1, Akt and GSK3β in TPN-fed mice, whereas treatment of chow-fed mice with an AhR inhibitor (CH223191) mimicked the effects of TPN. Thus, our results show that the gut microbiota and its metabolites induce insulin resistance, which are consistent with those reported by Natividad et al.(25). This study firstly revealed that TPN-induced insulin resistance is mediated by an alteration in the gut microbiota and a resulting decrease in the levels of AhR activators such as IAA and other indole derivatives.

Despite evidence linking the AhR to glucose metabolism and insulin sensitivity, the underlying mechanisms have yet not to be elucidated. It has been suggested that activation of the AhR improves insulin sensitivity and metabolic syndrome by enhancing the production of fibroblast growth factor-21(37) or intestinal interleukin-22.(26) However, the role of the AhR in metabolism is complex, and AhR activation has also been reported to be associated with insulin resistance.(38–40) Moreover, the AhR can ameliorate hepatic steatosis when activated by a plant extract(26) but aggravate hepatic steatosis when activated by environmental dioxin.(40) Therefore, the physiological and/or pathological processes downstream of AhR activation may vary between different organs or cells and between different types of ligand. The present study demonstrated that the plasma GLP-1 concentration was significantly lower in H-PN patients than in L-PN patients. Furthermore, an AhR inhibitor (CH223191) decreased the plasma GLP-1 level in chow-fed mice, while an AhR activator (Ficz) increased the GLP-l level in TPN-fed mice. GLP-1 exerts various beneficial metabolic effects including enhanced insulin secretion, improved insulin sensitivity, reduced appetite and delayed gastric emptying.(41) GLP-1 deficiency during TPN would be expected to have a detrimental effect on insulin sensitivity and ultimately result in metabolic disorders. According to our data, the GLP-1 analog, liraglutide was able to reverse the detrimental effects of TPN on insulin sensitivity and glycogen deposition in mice. We also found that liraglutide alleviated skeletal muscle atrophy and reduced visceral fat accumulation in mice during TPN. Skeletal muscle is the central organ for insulin-dependent glucose uptake and promotes insulin sensitivity,(42) while adipose tissue, especially visceral fat accumulation, is a key source of inflammatory insulin resistance.(43) Moreover, the preservation and accrual of lean mass is an important goal of TPN and an important predictor of clinical outcomes in patients with IF.(28, 44, 45) Our findings raise the possibility that GLP-1 might be a potential target for treating TPN-related metabolic complications in patients.

Enterogenous LPS leakage results in a mild chronic inflammatory condition that is a cause of insulin resistance.(46) LPS leakage from an impaired gut barrier during TPN is also related to clinical complications such as enterogenous infection or sepsis. Our observations showed that *Lactobacillus* abundance was negatively correlated with serum LPS concentration and that treatment with IAA reduced the circulating level of LPS in TPN mice. In view of these findings, additional research is merited to establish whether treatment with IAA might reduce the infectious morbidity and prevent metabolic complications in IF patients receiving long-term TPN.

In conclusion, this study shows for the first time that TPN induces glucose metabolism disorders through gut microbiota dysbiosis (including a decrease in the abundance of *Lactobacillaceae*). TPN decreases the levels of tryptophan metabolites (such as indole derivatives) by altering the gut microbiota, thereby inhibiting indole/AhR signaling and GLP-1 production, which leads to glucose metabolism disorders. Treatment with IAA or GLP-1 effectively prevents TPN-induced glucose metabolism disorders. This study highlights that GLP-1 supplementation and manipulation of the gut microbiota and its metabolites might represent effective strategies to prevent TPN-induced glucose metabolism disorders and their complications.

## Materials and Methods

### Clinical participants, sample collections and test

This retrospective analysis of a prospective database included patients diagnosed with IF who received continuous or intermittent PN for ≥2 weeks at the Clinical Nutrition Center of Jinling Hospital, Nanjing, Jiangsu, China. The main cohort of 256 patients was enrolled between August 2013 and August 2018 (Fig. S1). A secondary cohort of 16 patients (matched for age, intestinal anatomy and small intestinal length) was enrolled between 2018 and 2019 for analysis of the effects of PN on the intestinal flora. The inclusion criteria were: (1) aged 18–70 years old; (2) diagnosis of type 2 IF; (3) continuous or intermittent PN was given for ≥14 days; and (4) average energy provision by PN was ≥10% of requirements. The exclusion criteria were: (1) history of diabetes mellitus; (2) history of hepatic or renal insufficiency; and (3) pregnancy.

#### Nutritional strategy

The individualized nutritional schemes were adjusted daily according to resting energy expenditure (Quark PFT Ergo pulmonary function testing system, Cosmed, Rome, Italy), nitrogen balance, serum nutritional markers and fluid balance during hospitalization. The PN formula was mainly composed of dextrose, medium- and long-chain lipid emulsions, compound amino acids, fat-soluble and water-soluble vitamins, and functional nutritional elements such as fish oil, branched-chain amino acids, glutamine, arginine and albumin according to the specific needs of the patient.

#### Blood tests

For all 256 patients in the main cohort, fasting blood samples were drawn at 6:00 am on the second day after admission and then every 7 days during hospitalization. PN and enteral feeding were stopped overnight for measurement of fasting glucose and fasting insulin levels and blood biochemistry investigations (7600 Automatic Biochemical Analyzer, Hitachi, Tokyo, Japan). HOMA-IR was calculated as fasting glucose (mmol/L) × fasting insulin (mIU/L) / 22.5 (47). Sixty-nine of the 256 patients in the main cohort agreed to provide 2-mL blood samples during hospitalization for the measurement of GLP-1 and IAA levels. After isolation of serum by centrifugation (4°C, 4000×*g*, 15 min), the serum levels of total GLP-1 (ELISA kit, Merck, Darmstadt, Germany) and IAA (ELISA kit, MyBiosource, San Diego, CA, USA) were measured following standard protocols.

#### Assessment of clinical complications

Information regarding clinical complications in the 256 patients in the main cohort were extracted from the electronic medical records. The clinical complications included ICU transfer, emerging liver injury (both ALT and AST 200% above the upper limit of the normal range; AST > 40 U/L and AST/ALT ratio > 1; γ-glutamyl transpeptidase (γ-GT) > 200 U/L; or ultrasound or biopsy evidence of hepatic steatosis, necrosis and/or fibrosis), renal insufficiency (serum creatinine > 178 μmol/L), cardiovascular events (angina pectoris or myocardial infarction), fever, catheter-related infection, sepsis, pneumonia, and other infections during hospitalization.

#### Body composition

Body composition was evaluated using an S10 bioelectric impedance analyzer (InBody Co, Ltd., Seoul, Korea), which has a tetrapolar eight-point tactile electrode system and six different frequencies (1 kHz, 5 kHz, 50 kHz, 250 kHz, 500 kHz and 1000 kHz). Body composition measurements were made on the same mornings as the blood tests. Body protein mass proportion, fat-free mass proportion and skeletal muscle mass proportion were defined as body protein mass, fat-free mass and skeletal muscle mass relative to body weight, respectively.

#### Animals

C57BL/6J male mice weighing 25–30 g were randomly assigned to the chow group (*ad libitum* access to food and infusion of saline) and TPN group (fasted and given TPN) for 7 days. The TPN formula is shown in Table S1. Mice were kept on a 12/12 hour light/dark cycle. At the experimental endpoints, chow and TPN were withdrawn for 6 hours prior to collection of fasting blood samples and performance of intraperitoneal glucose tolerance tests (IPGTTs) and intraperitoneal insulin tolerance tests (IPITTs). Subsequently, the mice were sacrificed under anesthesia, and their serum and tissues were harvested. Stool samples were collected for bacterial 16S rRNA gene sequencing and used in oral gavage experiments.

The mice were divided randomly into four groups (n=3 for each group): Control, Abx, TPN→Abx and Chow→Abx. Mice in the Control group were given distilled water without antibiotics, while mice in the other groups were treated with a mixture of antibiotics (ampicillin 1 g/L, neomycin 1 g/L, metronidazole 1 g/L and vancomycin 500 mg/L) for 2 weeks to deplete the intestinal microbiota. Successful eradication of the intestinal microbiota was confirmed by analysis of fecal samples with qPCR. All mice were given ^60^Co-irradiated sterile food.[20] After being provided with sterile drinking water for 3 days, mice in the TPN→Abx and Chow→Abx groups were gavaged twice each week for 4 weeks with fresh cecal samples collected from mice fed with chow or TPN. The cecal samples were dissolved in sterile phosphate-buffered saline (PBS; Life Technologies, Carlsbad, CA, USA) at a concentration of 1 g per 5 mL, and 0.25 mL of the diluted sample was administered at each gavage.

#### Mouse model of TPN

Mice were housed five-to-a-cage and maintained in a climate-controlled facility with a 12:12-hour light-dark cycle for a period of 4 days before the surgical placement of a central venous catheter (CVC; silastic tubing with an internal diameter of 0.012 inches; Dow Corning, Midland, MI, USA) into the right jugular vein under pentobarbital anesthesia. The proximal end of the CVC was tunneled subcutaneously to exit at the tail. After surgery, each mouse was housed individually in a cage, and the CVC was connected to an infusion pump (48). The mice in both groups were allowed to eat and drink freely for 1 day after surgery and then received either chow or TPN (0.40 mL/h) for 7 days.

#### IPGGT and IPITT

Mice were fasted for 6 hours and then injected intraperitoneally with glucose (2.0 g/kg) for glucose tolerance tests or insulin (0.5 U/kg) for insulin tolerance tests. Blood glucose was measured at 0, 15, 30, 60, 90 and 120 min with a glucometer (Bayer, Leverkusen, Germany).

#### Measurement of serum indices

Blood samples were collected from the retro-orbital plexus of each mouse after anesthesia. Serum was acquired by centrifugation (4°C, 4000×*g*, 15 min) and frozen in liquid N_2_ until further analysis. Measurements of LPS (LPS ELISA kit, Cusabio, Wuhan, Hubei, China), IAA (IAA ELISA kit, MyBiosource, San Diego, CA, USA), GLP-1 (GLP-1 total ELISA kit, Merck, Darmstadt, Germany) and biochemical parameters including ALT, AST, ALP, total bilirubin, total protein and albumin (7600 Automatic Biochemical Analyzer, Hitachi, Tokyo, Japan) were performed using serum from nonfasted animals after 7 days of the nutritional intervention. Fasting glucose level (measured with a glucometer; Bayer, Leverkusen, Germany) and fasting insulin level (insulin mouse ELISA kit, Thermo Fisher Scientific, Waltham, MA, USA) were measured in fasted mice.

#### Liver histology and PAS staining

Samples of liver tissue from the left lobe were fixed in 4% paraformaldehyde solution overnight at 4°C. The fixed liver tissue was prepared as paraffin sections (5 µm thick) after dehydration with alcohol and stained with a PAS kit (Solarbio Life Science, Beijing, China).

#### Liver glycogen quantification

Liver tissue was collected, shredded and frozen in liquid N_2_. Quantification of liver glycogen in frozen liver tissue was carried out using a glycogen quantification kit (Solarbio Life Science, Beijing, China).

#### Immunoblotting

Total protein was extracted from the frozen liver tissue. Protein samples were separated by SDS-polyacrylamide gels and transferred to polyvinylidene fluoride gels. Antibodies against p-IRS1 (Ser307) (#2381), IRS1 (#2382), p-Akt (Ser473) (#4060), Akt (#9272), p-GSK3β (Ser9) (# 37F11), GSK3β (#5676), GAPDH (#2118) and β-actin (#4970) were used (1:1000 dilution; Cell Signaling Technology, Danvers, MA, USA). The expression of GLP-1 in the terminal ileum (within 10 cm of the cecum) and colon was examined with an anti-GLP-1 antibody (1:2000; #ab180443, Abcam, Cambridge, UK)

#### Quantitative RT-PCR to detect expression of the proglucagon gene

Total RNA was extracted from segments of the terminal ileum and colon and purified using TRIzol reagent (Takara Bio, Shiga, Japan). RNA was reverse transcribed into cDNA with PrimeScriptTM RT Master Mix (Takara Bio, Shiga, Japan). Quantitative RT-PCR was performed with SYBR® Premix Ex Taq™ (Takara Bio, Shiga, Japan) in a 7300 Real-Time PCR system (Applied Biosystems, Foster City, CA, USA). The mRNA expression of Gcg was calculated using the 2−ΔΔCT quantification method and normalized to that of the housekeeping gene, Gapdh. The primer sequences were as follows: Gcg, forward 5ʹ-ACAGCAAATACCTGGACTCC-3ʹ and reverse 5ʹ-CAATGTTGTTCCGGTTCCTC-3ʹ; Gapdh, forward 5ʹ-TGGCAAAGTGGAGATTGTTGCC-3ʹ and reverse 5ʹ-AAGATGGTGATGGGCTTCCCG-3ʹ.

#### Immunohistochemical staining of GLP-1

Small segments of the terminal ileum and colon were fixed, embedded in paraffin, dehydrated and incubated with an anti-GLP-1 antibody (1:800; Abcam, Cambridge, UK) overnight at 4°C after antigen retrieval and blocking for 1 hour. Then, the sections were processed with diaminobenzidine and counterstained with hematoxylin, and the coverslips were fixed with 50% neutral resins.

#### In vivo administration of drugs

IAA injections in mice were performed as described previously (49, 50). A stock solution of IAA (10 mg/mL; #87-51-4; Psaitong, Beijing, China) in PBS was prepared by adding NaOH (1 N) until the IAA had fully dissolved and then adjusting the pH to 7.4 with 25% (v/v) HCl. 6-formylindolo(3,2-b)carbazole (Ficz; 1 µg/day; Enzo Life Sciences, Farmingdale, NY, USA) and CH223191 (0.25 mg/day; MedChemExpress, Monmouth Junction, NJ, USA) were injected intraperitoneally to activate and inhibit the AhR, respectively.

Control animals were injected with vehicle (dimethyl sulfoxide [DMSO]). Liraglutide (0.3 mg/kg/day; Victoza, Novo Nordisk, Bagsværd, Denmark) or vehicle (normal saline) was injected subcutaneously.

### 16S rRNA amplicon sequencing and untargeted metabolomics

The cecal contents from TPN/Chow mice and fecal from type 2 IF patients were extracted under aseptic conditions and frozen in liquid N_2_ for characterization of the gut microbiota via 16S amplicon sequencing.

For 16S rRNA amplicon sequencing, DNA was extracted from the cecal contents and identified. One or several variable regions were selected, and universal primers were designed based on the conserved regions for PCR amplification. Then, the hypervariable regions were sequenced and identified. A small-fragment library was constructed by single-end sequencing using the IonS5^TM^XL sequencing platform. The abundance, diversity and composition of the gut microbiota were analyzed by read-shear filtration, operational taxonomic unit (OTU) clustering and species classification analysis. OTU cluster analysis was performed using Uparse 7.0.1001 (http://drive5.com/uparse/) (51). The heatmap was drawn using R 2.15.3 (https://www.R-project.org/). Calculation of the UniFrac distance and construction of the UPGMA sample clustering tree were carried out using Qiime 1.9.1 (52). The PCoA diagram was drawn in R 2.15.3, and linear discriminant analysis of effect size was performed using LEfSe.

The cecal contents from TPN/Chow mice were performed untargeted metabolomics. 1.0 mg of each sample was placed in a microcentrifuge tube, and 400 μL of 80% methanol solution was added. The tube was vortexed and centrifuged (14000×*g*) at 4°C for 20 min after standing for 60 min at −20°C. The supernatant was transferred to a 1.5-mL centrifuge tube and vacuum freeze-dried, and the residue was dissolved in 100 µL of complex solvent, vortexed again, and centrifuged (14000×*g*) at 4°C for 15 min. An equal amount of supernatant was transferred to high-performance liquid chromatography glass vials for untargeted liquid chromatography-mass spectrometry (LC-MS) analysis. Differential metabolites and KEGG pathways were obtained by data analysis. Quantification of metabolites was carried out using Compound Discoverer software (Thermo Fisher Scientific, Waltham, MA, USA). Charts were constructed using R 3.1.3.

## Supporting information

Supplement meterials

## Statistical analysis

No statistical methods were used to predetermine the sample size. All statistical analyses of clinical data were performed using SAS software, version 9.4 (SAS Institute, Cary, NC, USA). Experimental data were analyzed using Prism 5 (GraphPad Software, San Diego, CA, USA). Normally distributed data are presented as the mean ± standard error of the mean (SEM) and were compared between groups using the Student’s t-test for unpaired samples. Pearson correlation analysis was carried out to estimate the correlation between two numeric variables. Binary logistic regression was used to evaluate the association between HOMA-IR values and the risk of clinical complications. Two-tailed P-values <0.05 were considered indicative of statistical significance. For the differential gut microbiota and metabolites, the significance level was set at a variable importance in projection (VIP) score > 2.0, fold change > 2.0 or < 0.5, and P < 0.05.

## Data availability

The datasets used and/or analysed during the current study are available from the corresponding author on reasonable request.

## Ethics approval

All participants had provided informed written consent for their clinical data and blood samples (when provided) to be used in future research studies The clinical study received approval from the the ethics committee of Jinling Hospital approved the study (2014ZFYJ-010). All in vivo assays were performed in triplicate, and approved by The animal ethics committee of Jinling Hospital and also followed ARRIVE guidelines.

## Author Contributions

PW and HS takes responsibility for the content of the manuscript, including the data and analysis; Study concept and design: XW, PW and HS; The data acquisition: JY, GM, LZ, XG, YZ; Drafting of the manuscript: PW, HS, BX, CL, XW; Critical revision of the manuscript for important intellectual content: all authors; Statistical analysis: PW and HS; The order of the 2 co–first authors was assigned following the contribution of this ariticle. Study supervision: JL and XW. All authors read and approved the final manuscript.

## Acknowledgments

We thank all the patients who donated specimens for this study. This project was supported by the National Natural Science Foundation of China (81470797, 81770531), the Science Foundation of Outstanding Youth in Jiangsu Province (BK20170009), the National Science and Technology Research Funding for Public Welfare Medical Projects (201502022) and “The 13th Five-Year Plan” Foundation of Jiangsu Province for Medical Key Talents (ZDRCA2016091).

## Supplementary Figures

**Figure S1. Flow-chart summarizing the enrollment of the study participants.**

**Figure S2. Poor insulin sensitivity in mice given total parenteral nutrition (TPN). A–B.** Area under the curve values for intraperitoneal glucose tolerance tests (**A**) and intraperitoneal insulin tolerance tests (**B**). Mice received total parenteral nutrition (TPN group) or enteral feeding (Chow group) for 7 consecutive days. **C.** Venn diagram showing the potential central role of insulin resistance in glucose dysmetabolism, lean mass loss and clinical complications during parenteral nutrition.

**Figure S3. Fecal microbiota transplantation. A.** Heatmap showing the top 10 phyla in mice from the TPN and Chow groups. **B.** Heatmap showing the top 10 genera in mice from the TPN and Chow groups. **C.** Fold change in the fecal 16S rDNA content of mice measured using qPCR. **D–E.** Area under the curve values for intraperitoneal glucose tolerance tests (**D**) and intraperitoneal insulin tolerance tests (**E**).

**Figure S4. Levels of indole derivatives may underlie the association of reduced *Lactobacillus* abundance with glucose dysmetabolism. A.** Heatmap showing the top 15 differential metabolites. **B.** The top 10 most enriched Kyoto Encyclopedia of Genes and Genomes (KEGG) pathways for the differential metabolites. **C–D.** Area under the curve values for intraperitoneal glucose tolerance tests (**C**) and intraperitoneal insulin tolerance tests (**D**).

**Figure S5. Effects of aryl hydrocarbon receptor (AhR) activation and inactivation on glucose tolerance and insulin sensitivity. A–B.** Area under the curve values for intraperitoneal glucose tolerance tests (**A**) and intraperitoneal insulin tolerance tests (**B**) showing the effects of an inhibitor of the AhR (CH223191). **C–D.** Area under the curve values for intraperitoneal glucose tolerance tests (**C**) and intraperitoneal insulin tolerance tests (**D**) showing the effects of an agonist of the AhR (CH223191).

**Figure S6. Role of reduced glucagon-like peptide-1 production in parenteral nutrition-related impairment of insulin sensitivity. A.** Expression of *Gcg* mRNA in the terminal ileum and colon. **B.** Western blot analysis of GLP-1 expression in the terminal ileum and colon. **C–D.** Area under the curve values for intraperitoneal glucose tolerance tests (**C**) and intraperitoneal insulin tolerance tests (**D**). **E.** Gastrocnemius/body weight ratio. **F.** Epididymal fat/body weight ratio. **G.** Serum alanine transaminase (ALT) and aspartate transaminase (AST) concentrations. **H.** Liver edema evaluated from the liver/body weight ratio. **I.** Survival curves.

