## Supplement meterials for "Total parenteral nutrition drives glucose metabolism disorders by modulating gut microbiota and its metabolites"

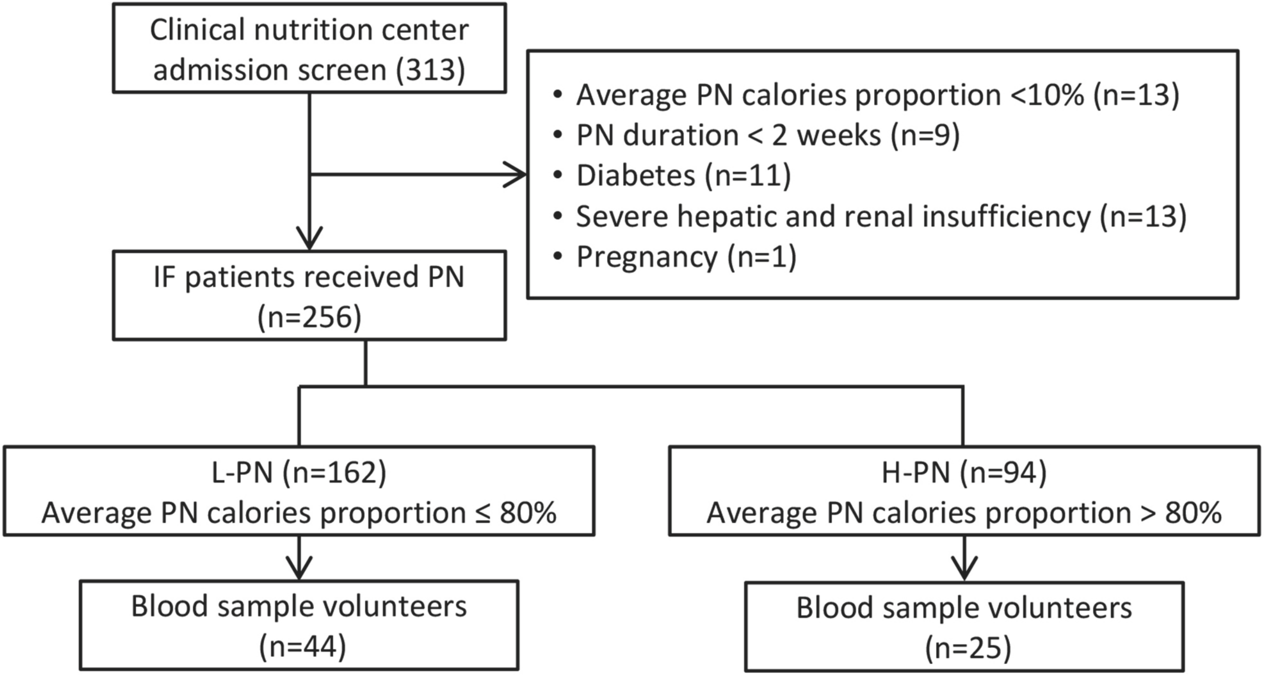


Figure S1. Flow-chart summarizing the enrollment of the study participants.

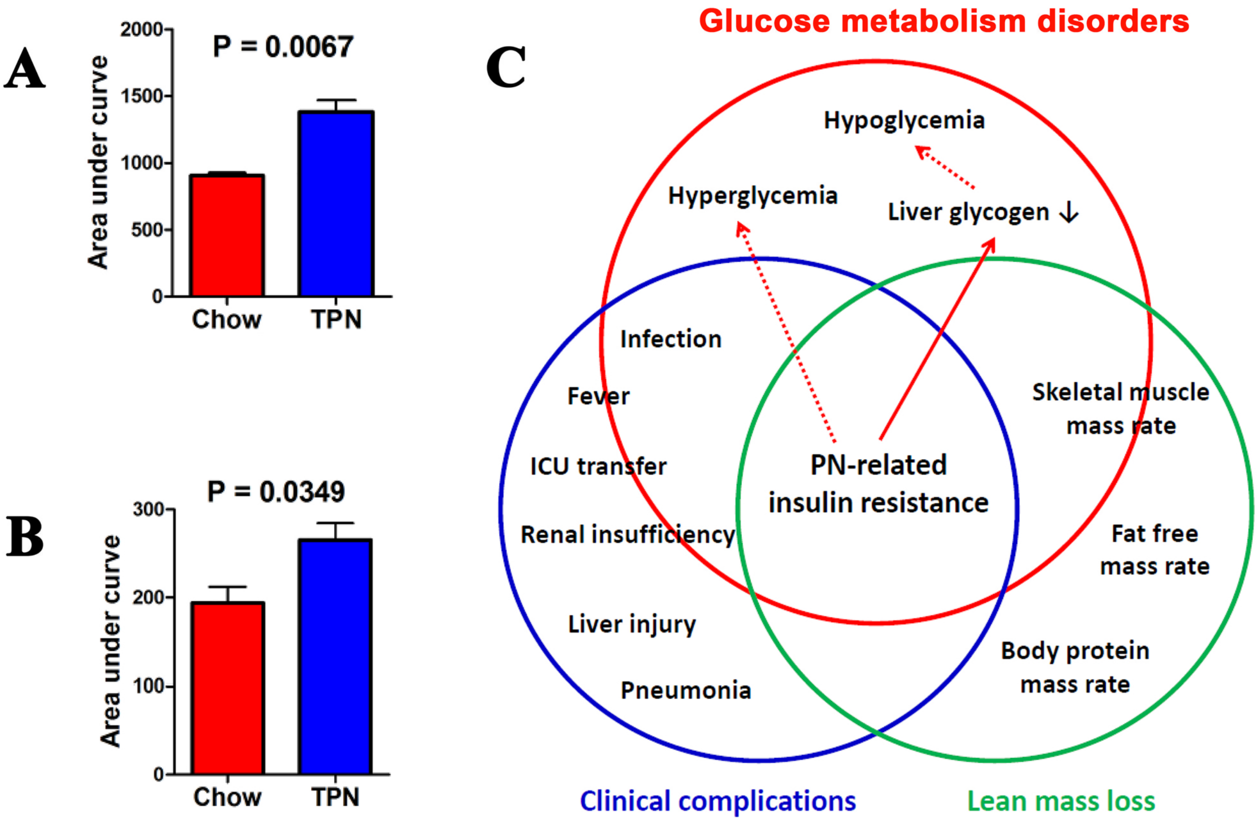


Figure S2. Poor insulin sensitivity in mice given total parenteral nutrition (TPN).

A–B. Area under the curve values for intraperitoneal glucose tolerance tests (A) and intraperitoneal insulin tolerance tests (B). Mice received total parenteral nutrition (TPN group) or enteral feeding (Chow group) for 7 consecutive days.

C. Venn diagram showing the potential central role of insulin resistance in glucose metabolism disorders, lean mass loss and clinical complications during parenteral nutrition.

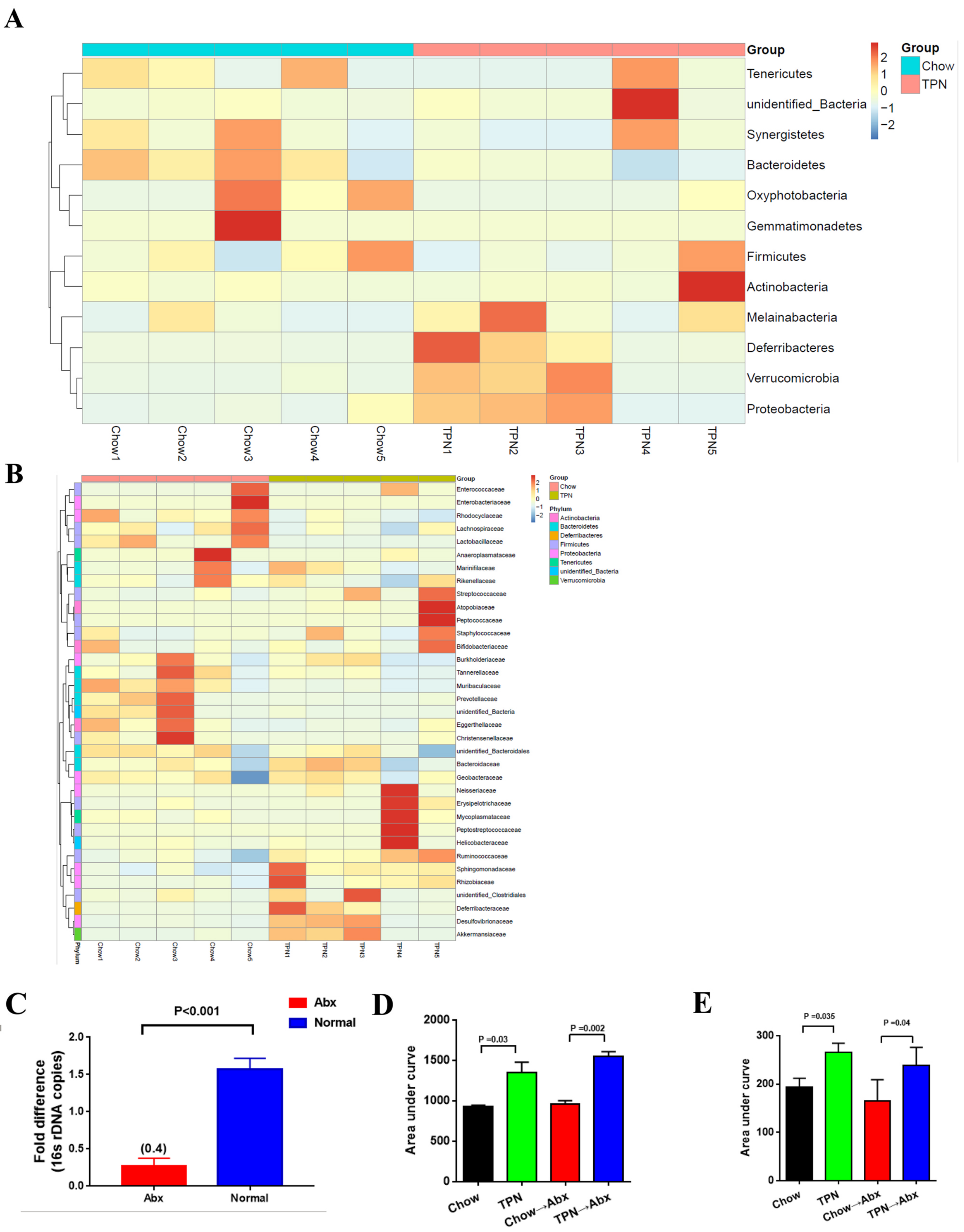


Figure S3. Fecal microbiota transplantation.

A. Heatmap showing the top 10 phyla in mice from the TPN and Chow groups.

B. Heatmap showing the top 10 genera in mice from the TPN and Chow groups.

C. Fold change in the fecal 16S rDNA content of mice measured using qPCR.

D–E. Area under the curve values for intraperitoneal glucose tolerance tests (D) and intraperitoneal insulin tolerance tests (E).

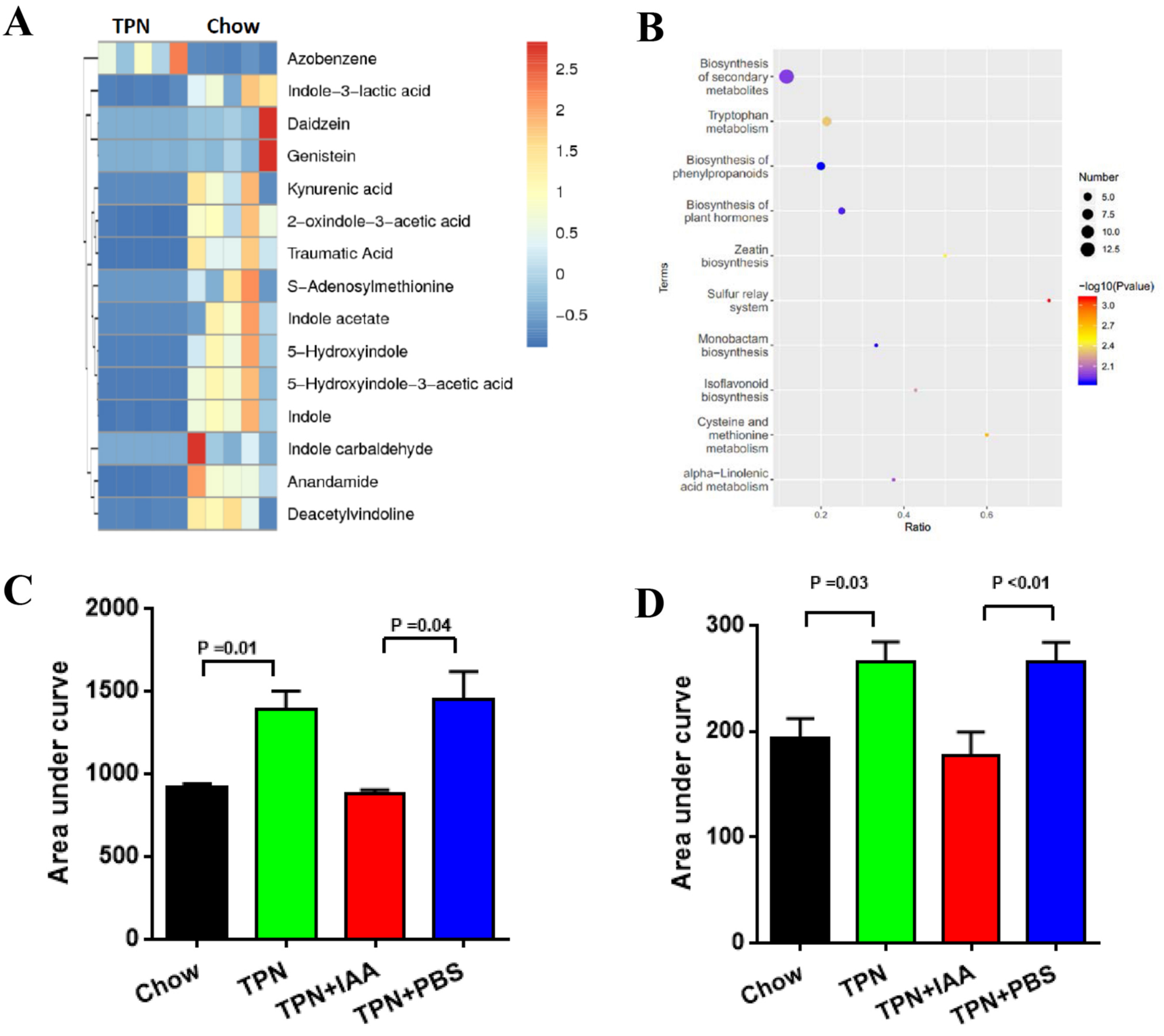


Figure S4. Levels of indole derivatives may underlie the association of reduced Lactobacillus abundance with glucose metabolism disorders.

A. Heatmap showing the top 15 differential metabolites.

B. The top 10 most enriched Kyoto Encyclopedia of Genes and Genomes (KEGG) pathways for the differential metabolites.

C–D. Area under the curve values for intraperitoneal glucose tolerance tests (C) and intraperitoneal insulin tolerance tests (D).

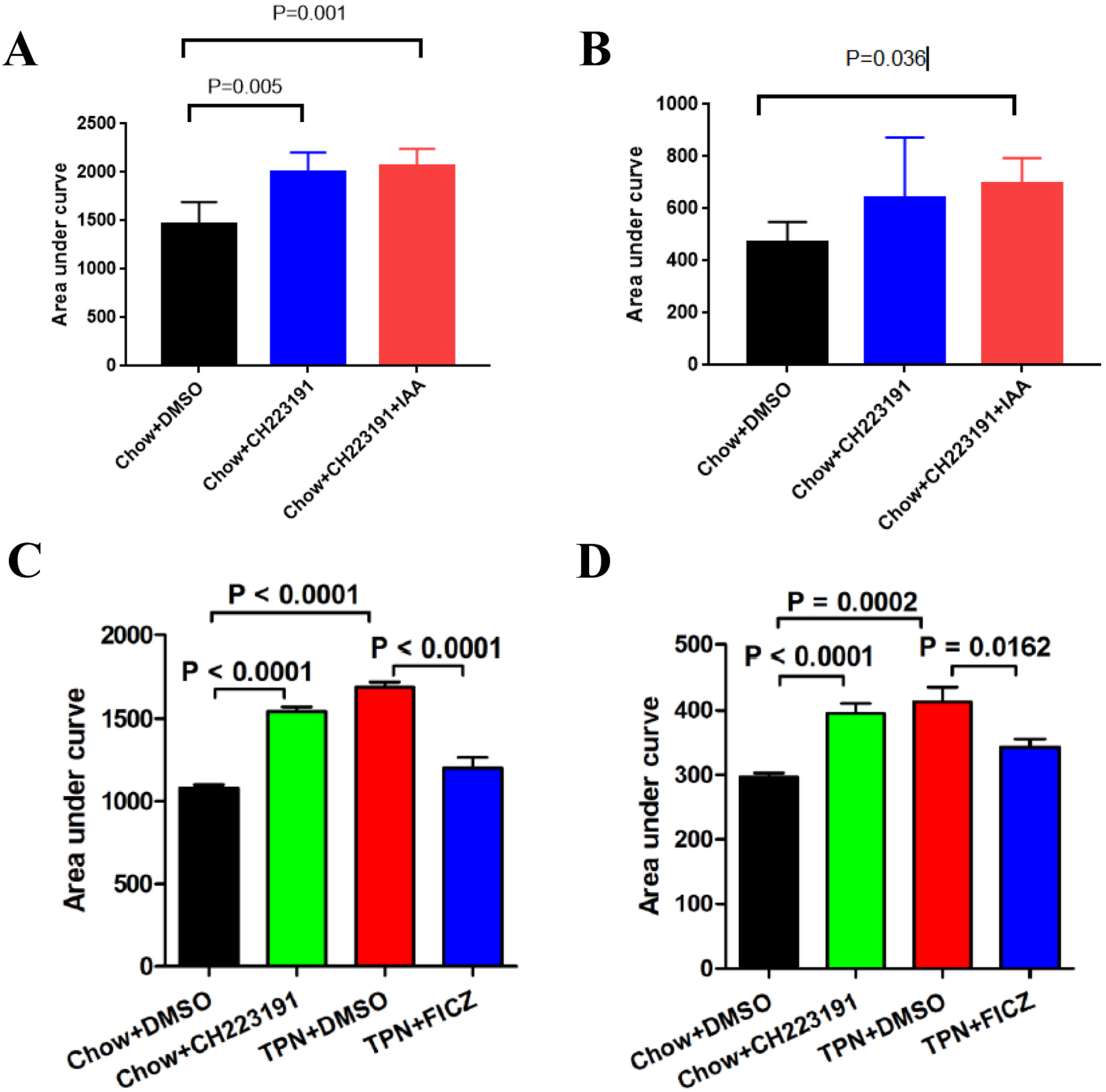


Figure S5. Effects of aryl hydrocarbon receptor (AhR) activation and inactivation on glucose tolerance and insulin sensitivity.

A–B. Area under the curve values for intraperitoneal glucose tolerance tests (A) and intraperitoneal insulin tolerance tests (B) showing the effects of an inhibitor of the AhR (CH223191).

C–D. Area under the curve values for intraperitoneal glucose tolerance tests (C) and intraperitoneal insulin tolerance tests (D) showing the effects of an agonist of the AhR (CH223191).

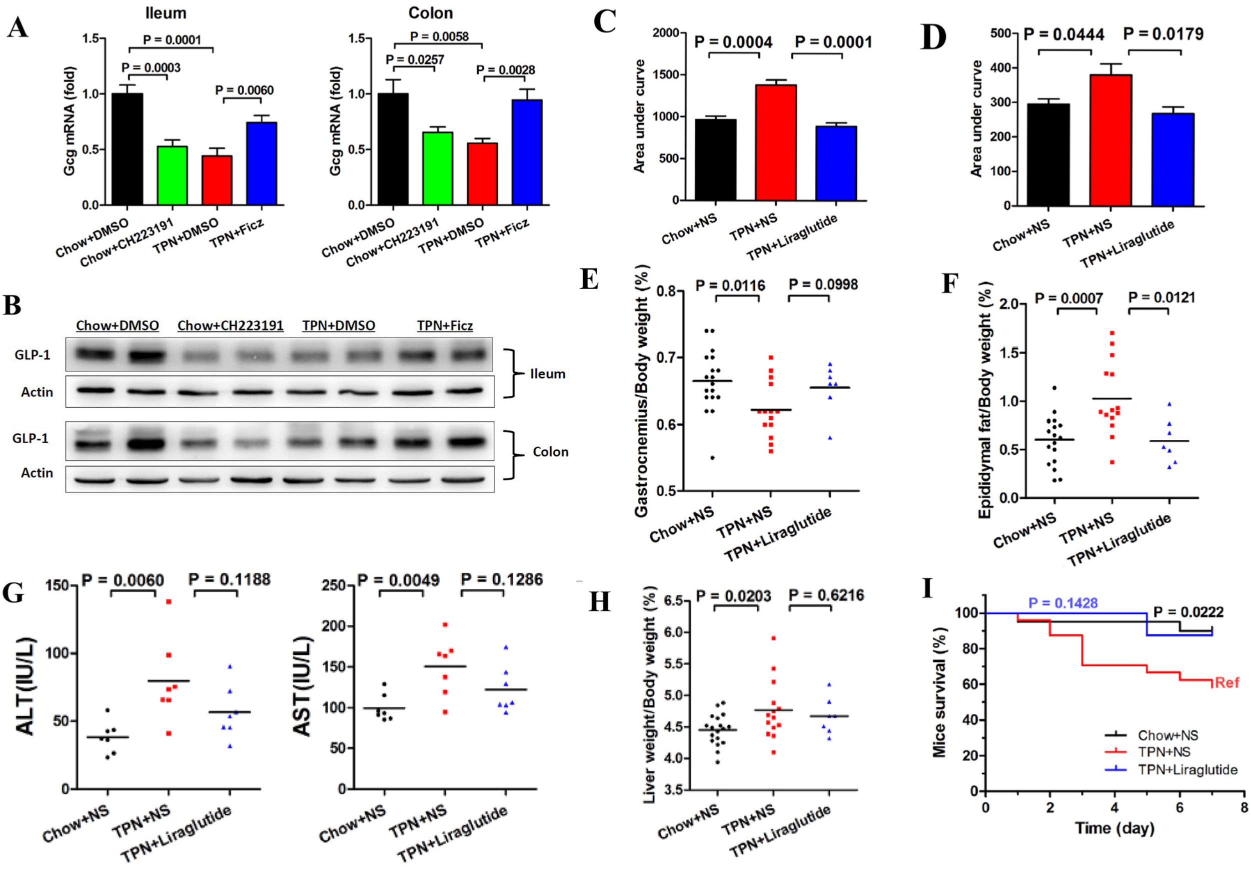


Figure S6. Role of reduced glucagon-like peptide-1 production in parenteral nutrition-related impairment of insulin sensitivity.

A. Expression of Gcg mRNA in the terminal ileum and colon.

B. Western blot analysis of GLP-1 expression in the terminal ileum and colon.

C–D. Area under the curve values for intraperitoneal glucose tolerance tests (C) and intraperitoneal insulin tolerance tests (D).

E. Gastrocnemius/body weight ratio.

F. Epididymal fat/body weight ratio.

G. Serum alanine transaminase (ALT) and aspartate transaminase (AST) concentrations.

H. Liver edema evaluated from the liver/body weight ratio.

I. Survival curves.

**Table S1.** Total parenteral nutrition composition (per 500 mL)

| Ingredient | Volume (mL) |
|---|---|
| 11.4% Compound amino acids | 220 |
| 80% Glucose | 167 |
| Intralipid (Fresenius Kabi) | 35 |
| 10% sodium chloride | 14 |
| 10% potassium chloride | 4 |
| 25% magnesium sulphate | 2 |
| 10% compound phosphate | 10 |
| 10% calcium gluconate | 12 |
| Microelements | 2 |
| Water-soluble vitamins | 8 |
| Fat-soluble vitamins | 8 |
| 50% potassium acetate | 8 |
| Normal saline | 10 |

**Table S2.** Parenteral nutrition-related characteristics for patients with intestinal failure

| Characteristic | n | | PN calorie proportion | | P value |
|---|---|---|---|---|---|
| | | | >80% | ≤80% | |
| N | 256 | | 94 (36.7%) | 162 (63.3%) | |
| PN duration, mean (range), days | 255 | 38.8 (14–208) | 41.4 (14–208) | 35.9 (20–208) | 0.026 |
| PN calorie proportion (%) | 249 | 59.43±1.84 | 91.2±0.75 | 41.1±1.56 | <0.001 |
| Basal metabolic rate, kcal/kg/day | 246 | 26.78±0.26 | 25.96±0.46 | 27.26±0.31 | 0.018 |
| Daily energy, kcal/kg | 200 | 29.10±0.48 | 27.98±0.88 | 29.75±0.56 | 0.077 |
| Daily energy via EN, kcal/kg | 202 | 14.66±0.81 | 4.72±0.62 | 20.4±0.89 | <0.001 |
| Daily energy via PN, kcal/kg | 202 | 17.61±0.64 | 23.49±0.85 | 14.21±0.73 | <0.001 |
| Small bowel length | 75 | 70.9±45.6 | 71.6±48.1 | 69.9±42.5 | 0.064 |
| Underlying pathophysiology, n (%) | 256 | | | | |
| Short bowel syndrome | | 75 (29.3%) | 36 (38.3%) | 39 (24.1%) | 0.016 |

| | | | | |
|---|---|---|---|---|
| Ileus | 60 (23.4%) | 30 (31.9%) | 30 (18.5%) | 0.015 |
| Extensive parenchymal disease | 54 (21.1%) | 13 (13.8%) | 41 (25.3%) | 0.030 |
| Intestinal dysmotility | 46 (18.0%) | 13 (13.8%) | 33 (20.4%) | 0.189 |
| Intestinal fistula | 3 (1.2%) | 0 (0%) | 3 (1.9%) | 0.184 |
| Other (including scleroderma) | 18 (7.0%) | 2 (2.1%) | 16 (9.9%) | 0.019 |

Values are presented as mean ± standard error of the mean unless otherwise stated. EN, enteral nutrition; PN, parenteral nutrition.

**Table S3.** Clinical characteristics of the 256 patients with intestinal failure

| Characteristic | PN calorie proportion | | P value |
|---|---|---|---|
| | >80% | ≤80% | |
| N | 94 (36.7%) | 162 (63.3%) | |
| Gender | | | 0.598 |
| Male | 53 (56.4%) | 98 (60.5%) | |
| Female | 41 (43.6%) | 64 (39.5%) | |
| Age, years | 47.7±1.49 | 44.3±1.28 | 0.102 |
| LOS, days | 41.2±3.20 | 25.4±0.90 | <0.001 |
| Hospitalization expenses, $10^3$￥ | 109.4±7.49 | 54.2±2.68 | <0.001 |
| Height, cm | 165.4±0.84 | 166.1±0.63 | 0.497 |
| BMI, kg/$m^2$ | 17.3±0.33 | 16.2±0.23 | 0.003 |
| NRS 2002 | 3.94±0.074 | 3.53±0.058 | <0.001 |
| PG-SGA | | | 0.001 |
| A grade | 1 (1.1%) | 9 (5.7%) | |
| B grade | 23 (26.4%) | 73 (46.5%) | |
| C grade | 63 (72.4%) | 75 (47.8%) | |
| Fasting blood glucose, mmol/L | 6.27±0.16 | 5.14±0.09 | <0.001 |
| Fasting blood insulin, mIU/L | 10.57±0.58 | 6.33±0.36 | <0.001 |
| HOMA-IR | 3.23±0.19 | 1.37±0.09 | <0.001 |
| Body protein mass proportion | 0.163±0.0016 | 0.171±0.0012 | <0.001 |
| Fat free mass rate proportion | 0.854±0.0083 | 0.887±0.0060 | 0.001 |

| | | | |
|---|---|---|---|
| Skeletal muscle mass proportion | 0.448±0.0046 | 0.470±0.0036 | <0.001 |

Values are presented as mean ± standard error of the mean or n (%). BMI, body mass index; HOMA-IR, homeostasis model assessment insulin resistance index; LOS, length of stay in hospital; NRS 2002, Nutritional Risk Screening 2002; PG-SGA, Patient-Generated Subjective Global Assessment; PN, parenteral nutrition.

Table S4. Clinical characteristics for 16 patients with intestinal failure.

| Group | No. | Age | Gender | LPS (EU/ml) | Pathophysiology | Length of SI (cm) | HOMA-IR * | Sample collection time (0 = first day of PN) | Calorie provision by PN (%) * | Sample type | Abundance of g-Lactobacillus (%) * |
|---|---|---|---|---|---|---|---|---|---|---|---|
| L-PN[a] | 1 | 67 | M | 0.040 | SBS-3 | 65 | 0.60 | Day-4 | 14 | Feces | 8.37 |
| | 2 | 46 | M | 0.053 | SBS-3 | 130 | 0.46 | Day-0 | 0 | Feces | 18.04 |
| | 3 | 39 | F | 0.120 | SBS-3 | 60 | 0.45 | Day-2 | 0 | Feces | 7.46 |
| | 4 | 32 | F | 0.020 | SBS-2 | 150 | 0.65 | Day-3 | 11 | Feces | 1.87 |
| | 5 | 60 | M | 0.038 | SBS-2 | 80 | 0.67 | Day-0 | 0 | Feces | 8.67 |
| | 6 | 63 | F | 0.047 | SBS-1 | 90 | 0.77 | Day-0 | 0 | Jejunostomy liquid | 49.74 |
| | 7 | 50 | M | 0.012 | SBS-1 | 60 | 0.24 | Day-1 | 0 | Jejunostomy liquid | 40.09 |
| | 8 | 48 | M | 0.019 | SBS-2 | 15 | 0.03 | Day-8 | 12 | Feces | 42.91 |
| | 9 | 38 | F | 0.056 | Vascular disease | - | 0.60 | Day-0 | 0 | Feces | 1.12 |
| H-PN[b] | 10 | 55 | M | 1.269 | SBS-2 | 70 | 5.89 | Day-18 | 91 | Feces | 1.74 |
| | 11 | 40 | F | 2.779 | SBS-3 | 60 | 2.30 | Day-49 | 83 | Feces | 0.14 |
| | 12 | 64 | F | 2.345 | SBS-2 | 90 | 9.84 | Day-46 | 87 | Feces | 1.16 |
| | 13 | 18 | M | 1.098 | SBS-3 | 15 | 4.79 | Day-19 | 89 | Feces | 2.27 |
| | 14 | 31 | F | 0.067 | SBS-1 | 150 | 8.10 | Day-11 | 100 | Jejunostomy liquid | 0.30 |
| | 15 | 35 | M | 0.085 | Vascular disease | - | 2.40 | Day-15 | 83 | Feces | 1.58 |
| | 16 | 61 | M | 1.260 | SBS-3 | 90 | 4.10 | Day-14 | 85 | Feces | 12.46 |

[a] PN accounting for ≤80% of total calories; [b] PN accounting for >80% of total calories.

HOMA-IR: Homeostasis Model Assessment Insulin Resistance index; LPS, lipopolysaccharides; PN: parenteral nutrition; SBS: short bowel syndrome; SI: small intestine.

P-values for the variables: length of SI, P = 0.931; pathophysiology, P = 0.618; age, P = 0.439; gender, P = 0.949; CRP: 0.224. * $P < 0.05$.
